# Enriching microlaser-labelled cells by commercial flow cytometry-based cell sorting systems

**DOI:** 10.64898/2026.10.02.756078

**Authors:** Matthias König, Tomasz T. Plaskocinski, Anna Popczyk, Vera M. Titze, Felix Gaedke, Maximilian Germer, Anna-Lena Schumacher, Annette M. Schmidt, Marcel Schubert, Malte C. Gather

**Affiliations:** Humboldt Centre for Nano- and Biophotonics, Institute for Light and Matter, Department of Chemistry and Biochemistry, University of Cologne, Cologne, Germany; Centre of Biophotonics, SUPA, School of Physics and Astronomy, University of St Andrews, St Andrews, UK; Faculty of Mathematics and Natural Sciences, Cluster of Excellence Cellular Stress Responses in Aging-associated Diseases (CECAD), University of Cologne, Cologne 50931, Germany; FACS & Imaging Core Facility, Max Planck Institute for Biology of Ageing, Cologne, Germany; Institute for Light and Matter, Department of Chemistry and Biochemistry, University of Cologne, Cologne, Germany

## Abstract

Microlasers based on whispering-gallery modes (WGMs) provide bright, spectrally narrow emission, making them ideal optical probes for high-resolution cellular tagging, single-cell tracking, and biosensing. To reproducibly apply WGM microlasers in biomedical or tissue-engineering contexts, control over the number of lasers in a cell population is highly desirable. Here we use a commercial instrument for flow cytometry-based cell sorting to obtain HEK293T cell populations with a controlled number of polystyrene laser particles (LPs). By modifying the particle surface with poly(ethyleneimine) (PEI) or lipofectamine, we enhance LP-association to cells. We demonstrate that our sorting procedure yields viable cells showing unchanged proliferation rates. Manufacturer-dependent differences in particle-cell interactions were observed and investigated by scanning and transmission electron microscopy (SEM/TEM) and flow cytometry. Finally, we show that the same sorting approach enables the generation of multi-cellular spheroids with a defined ratio of LP-containing and LP-free cells. This work lays the foundation for larger tissue engineering studies with defined LP populations and control over LP-placement with respect to the cell.

## Introduction

Over the last decade, spherical lasing particles (LPs) have emerged as powerful photonic tools to study biological processes, owing to their unique optical characteristics and compatibility with living systems. These lasers often are spherical particles made from a transparent polymer, usually polystyrene, that confine light as a whispering-gallery mode (WGM). When loaded with a highly fluorescent organic dye, these WGM LPs generate laser light under nanosecond optical excitation, with the pulse energies required for lasing often in the nanojoule range, well below the damage threshold of most biological tissue. The narrow and bright emission peaks of these LPs provide distinct advantages over conventional fluorescent markers and probes, enabling high-resolution signal discrimination and fast readout^1–4^. The emission spectrum of WGM LPs is determined by their diameter, the fluorescent dye within the polystyrene and the refractive index of their immediate environment. Their high sensitivity to refractive-index changes in the surrounding environment makes them excellent sensors, e.g. for detecting binding events of antibodies or DNA^5,6^ to the surface of functionalized LPs. Polystyrene LPs can be internalized by various cells, extending the refractive-index sensing to intracellular processes. This sensing has been used to measure cardiac contractility in isolated cardiomyocytes as well as live zebrafish^7^. Both phagocytic and non-phagocytic cells engulf LPs with diameters up to their own size, and uptake can further be improved by commercially available surface modifications^8,9^. Internalized LPs can also be used for multiplexed barcoding and tracking of cells in three-dimensional skin or cancer models^3,10,11^. A recent study extended the refractive index sensing to the uptake of LPs by reef building corals^12^. This proves that LPs can also be used as a model system to study the impact of environmental microplastic on human health *in vitro*. Although thousands of labelled cells can easily be identified in a cell culture dish, it is essential to control the purity and composition of the underpinning cell population in terms of LP labelling density to fully unlock advanced tissue-engineering techniques.

One approach for the physical separation and isolation of defined cell populations is flow cytometry-based cell sorting. Flow cytometry is used to identify and characterize cell populations based on intrinsic light-scattering properties and fluorescence signals from labelled cellular components, such as antigens, while the sorting module subsequently separates the selected cells into distinct fractions^13–15^. More recently, this approach has been extended beyond simple detection of fluorescence to custom-built microfluidic setups in which the sorting decision is based on the spectral barcodes of small LPs inside the cells^16,17^. This approach allowed multipass flow cytometry measurements of single cells and the immune phenotyping of 32 different proteins^18^. However, such spectral flow cytometry setups are complicated to build, maintain and operate. In contrast, commercial cell sorting systems are often available in core facilities and operated by designated technical staff.

In this study, we leverage the widespread availability of such commercial flow cytometry and cell sorting systems to produce purified populations of cells labelled with large LPs, without the need for custom microfluidic systems. The gating strategy used during cell sorting is based on the forward- and side-scattering properties of the labelled cells as well as the fluorescence intensity of a membrane impermeable nuclear stain (DAPI, to label cells with ruptured membrane, i.e. dead cells) and of the gain medium within the LPs themselves. We then use these purified populations to reveal pronounced, manufacturer-dependent differences in LP uptake and localization, which we characterize by electron microscopy. Furthermore, we show how flow cytometry-based cell sorting can be applied to precisely control the number of LPs per spheroid, which is of particular relevance for tissue-engineering and deep-tissue sensing.

## Results

### Fluorescence brightness and uniformity determine LP lasing performance

To test whether standard flow cytometric cell sorting devices can be used to purify microlaser-containing cell cultures, we first followed established protocols for the addition of LPs to cells, employing two types of LPs that were also used in previous works^7,8,10^. Cells were first incubated with LPs in a cell culture dish, then collected from the dish and finally sorted into populations containing cells with LPs (LP^+^), bare LPs, and cells without LPs (LP^−^), based on forward scatter area (FSC-A), side scatter area (SSC-A), and the photoluminescence intensity of the LPs gain medium (Red-PL). For low-throughput applications, such as electron microscopy, pure LP^+^ are seeded, whilst for tissue engineering applications LP^+^ and LP^-^ are mixed at a defined ratio and seeded directly into ultra-low attachment plates (Fig. 1A).

**Fig. 1.**
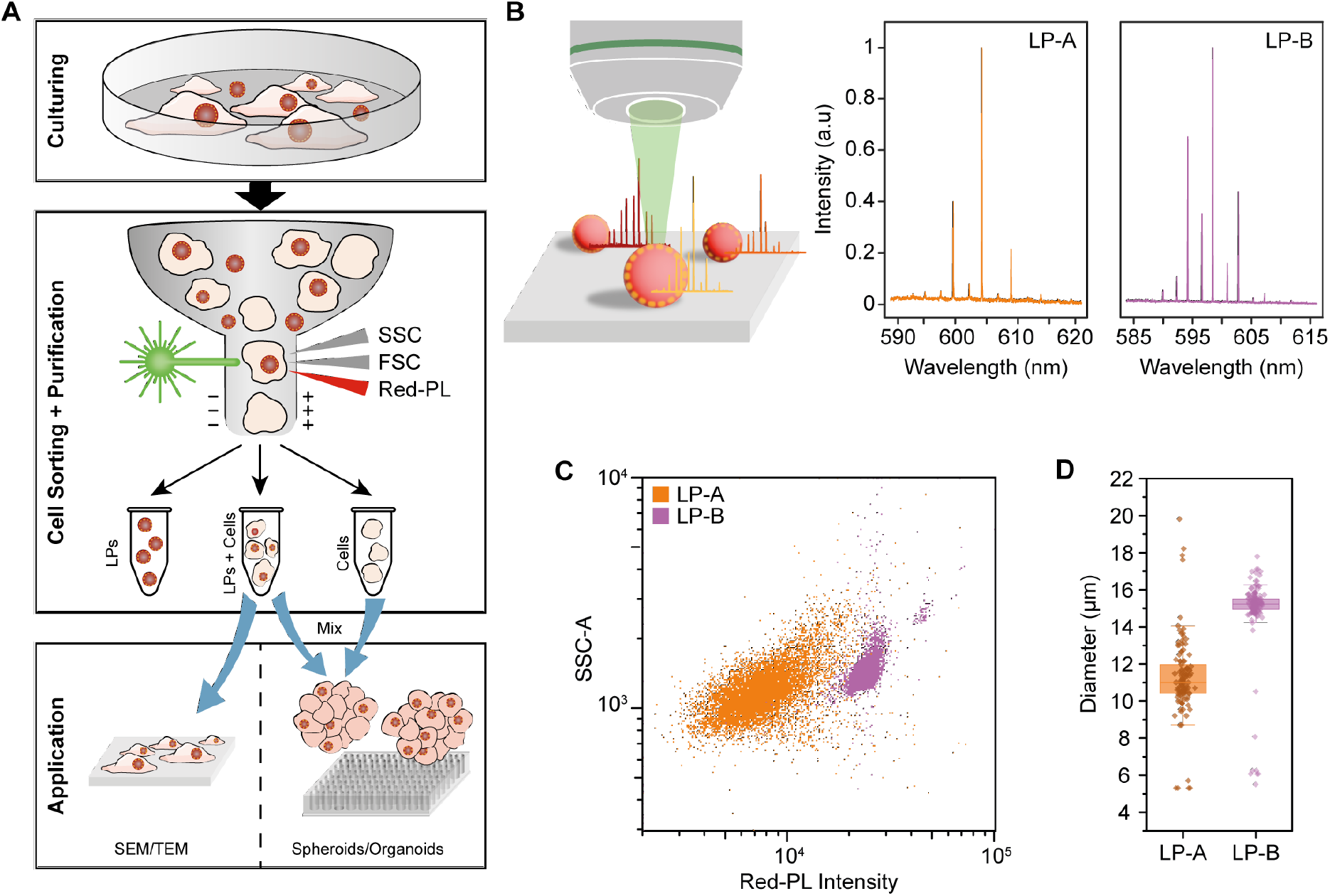
**A** Illustration of the purification and tissue-engineering workflow. Cells are first incubated with LPs and subsequently sorted based on their scattering properties. Next, cells are seeded for SEM/TEM analysis or mixed at defined ratios of LP-containing to unlabeled cells for spheroid formation. **B** LPs are excited by optical pumping (left); the spectral composition of the WGM emission depends primarily on the diameter of the LP and the fluorescent dye inside the polystyrene (PS) matrix (right). **C** Side scatter (SSC-A) signal versus red fluorescence intensity (Red-PL) for both LP-A and LP-B. *n=10,000*. **D** Average diameter of LP-A and LP-B. Box plots show the median and 25% and 75% quartiles; whiskers span all values that are not considered outliers. *n*_*LP-A*_ = *114, n*_*LP-B*_ = *138*.

To evaluate LP performance for hyperspectral applications, we first compared two specific LP types from two different suppliers (hereafter LP-A and LP-B) in the absence of cells. Both LP types showed characteristic WGM lasing peaks at around 600 nm (Fig. 1B). However, only 7 ± 2% of the LP-A sample showed lasing when pumped at 4 nJ, increasing to 40 ± 4 % at a pump pulse energy of 34 nJ. By contrast, all tested LP-B exhibited robust lasing already at 4 nJ (SI Fig. 1A).

To identify factors that may account for this difference in lasing performance, we investigated three LP properties: photoluminescence brightness (Red-PL, as a proxy for the amount of gain medium present in the polystyrene matrix), particle size, and particle uniformity. Flow cytometry revealed that LP-B were about 3.5 times brighter than LP-A (SI Fig. 1B). Additionally, LP-A showed a broader spread in the SSC-A vs Red-PL channel than LP-B (Fig. 1C), indicating a broader distribution in size and in the amount of gain medium contained in each particle. In contrast, LP-B showed two separated populations in the SSC-A vs Red-PL channel (Fig. 1C) as well as in FSC-A vs SSC-A plots (SI Fig. 2C), with the smaller population being of higher Red-PL intensity. The higher Red-PL intensity and stronger scattering properties of the smaller population can be attributed to agglomerates of LPs. We also performed brightfield microscopy of LPs to determine their size. This revealed that LP-A had an average diameter of 11.2 ± 2.1 µm, whereas LP-B averaged 14.9 ± 2.1 µm (Fig. 1D). We therefore conclude that a high, uniform brightness and a narrow distribution in forward and side scatter are favorable for reliable lasing.

### Surface modification alters cell-LP interaction

In addition to their optical and geometrical properties, LPs must adhere to or be internalized by the cells they are supposed to label. Adhesion and internalization are strongly influenced by the physical and chemical surface properties of the used LPs. To investigate the uptake of LPs, we chose HEK293T cells expressing EGFP in the membrane (HEK-EGFP) to enable detection of LP internalization by subsequent fluorescence microscopy. We incubated HEK-EGFP cells with LPs and, after washing the cells with DPBS, analyzed their interaction with the LPs using brightfield and fluorescence microscopy.

LP-A were primarily localized on or beside cells, whereas LP-B showed little to no attachment to cells (Fig. 2A). To enhance overall cell–LP interactions, LPs were coated with either lipofectamine or polyethyleneimine (PEI), thereby reversing the LP’s surface charge from negative to positive, which promotes interaction with the negatively charged cell membrane^19^ (Fig. 2B, SI Fig. 1C). The coating generally improved cell association, with PEI-coated LPs exhibiting the strongest interaction with cells, indicated by the localization of LPs exclusively on the cells after washing (Fig. 2A).

**Fig. 2.**
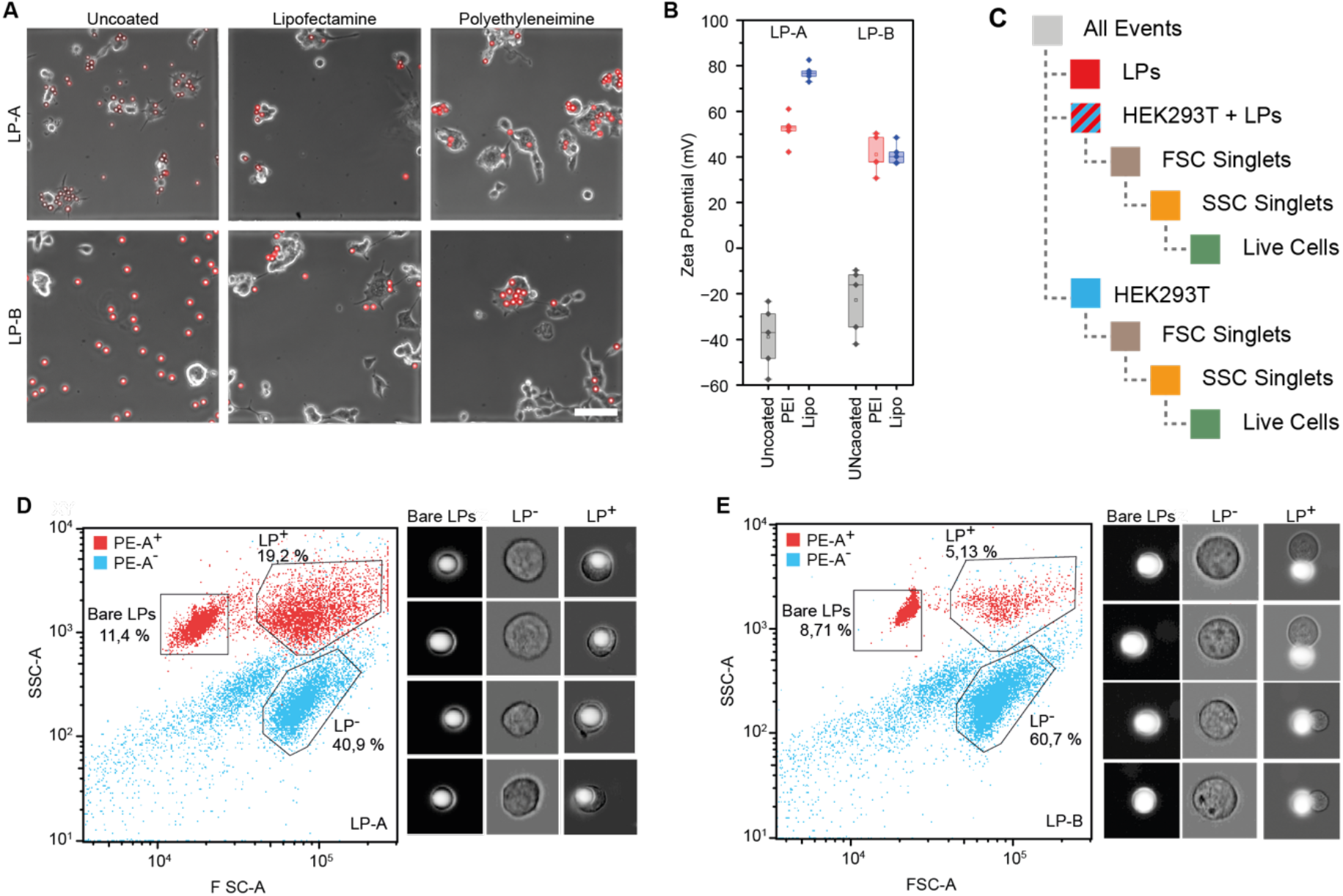
**A** Brightfield microscopy overlaid with fluorescence images of HEK293T cells incubated with LP-A and LP-B without any treatment (left), lipofectamine coating (center), and polyethyleneimine coating (right) after a wash with DPBS. Scale bar, 100 µm. **B** Surface charge of uncoated and coated LPs, for both LP types. **C** Gating strategy for HEK293T cells with LPs. **D** Scatter plots of HEK-EGFP cells incubated with LP-A, showing gates for bare LPs, LP^-^ cells, and LP^+^ cells. Events are colored by red LP fluorescence in the Red-PL channel, with fluorescent events indicated in red and non-fluorescent events indicated in blue. Brightfield images show representative events from an imaging cytometer for the corresponding previously sorted populations. **E** Same as D for HEK-EGFP cells incubated with LP-B.

Next, we sorted the cells incubated with PEI-coated LPs by flow cytometry. We followed a simple gating strategy that first identified the three major populations of LP^-^, LP^+^ and bare LPs based on their FSC-A and SSC-A characteristics. The LP^−^ and LP^+^ cell populations were subjected to singlet discrimination using FSC- and SSC-based parameters. Finally, viable singlet cells were selected by excluding DAPI-positive events and sorted accordingly. Because LP-containing cells were significantly enlarged, it was essential to use 130-µm nozzles and flow the samples at a pressure of 14 psi (Fig. 2C, SI Fig. 2A). For both LP types, LP^-^, LP^+^ and bare LP populations were clearly distinguishable in the flow cytometry scatter plots (Fig. 2D & E, SI Fig. 2B & C). For LP-A, 19.2% of events were LP^+^, compared with only 5.1% for LP-B, indicating reduced internalization efficiency for LP-B. Additional experiments with an imaging flow cytometer confirmed that the sorted populations consisted exclusively of the intended target groups. Analysis of the images also indicated that LP-B were attached to the cell surface, while LP-A seemed to be fully internalized. We also assessed the impact of LP labelling on the proliferation rate of HEK293T cells. Cells associated with LP-A as well as the LP^-^ fraction showed similar growth to the HEK-EGFP control group, whereas both LP^+^ and LP^-^fractions of cells incubated with LP-B showed slower growth resulting in 20% less confluency after 5 days in culture (SI Fig. 2E).

### Scanning and transmission electron microscopy reveals LP–cell interaction

We found that confocal microscopy could not always reliably resolve whether LPs had been fully engulfed by the cell, mostly because the interface between the cell membrane and the LP is often obscured by a lensing effect of the polystyrene LPs (SI Fig. 3A & B). To overcome this problem, sorted populations of cells carrying either LP-A or LP-B were seeded at single-cell densities for SEM and TEM analysis (Fig. 3A). SEM imaging of the purified cultures revealed that the cell membrane of HEK293T cells completely engulfed LP-A, whereas LP-B were positioned on top of the cells, with their smooth polystyrene surfaces clearly visible (Fig. 3B). To further investigate the cell–LP interface, thin sections of the cells were examined by TEM. LP-A were fully surrounded by cytosol, forming a distinct interface between the LP and the surrounding cytosol, with no separating membrane visible. In contrast, LP-B were only partially engulfed, with the cell membrane forming an outward-reaching cup that covered the cell-facing portion of the LP, while no direct contact between the LP and the cytosol was observed (Fig. 3C). SEM imaging further showed that cells sometimes interacted with several LPs simultaneously and at times even internalized several LP-A (SI Fig. 3C).

**Fig. 3.**
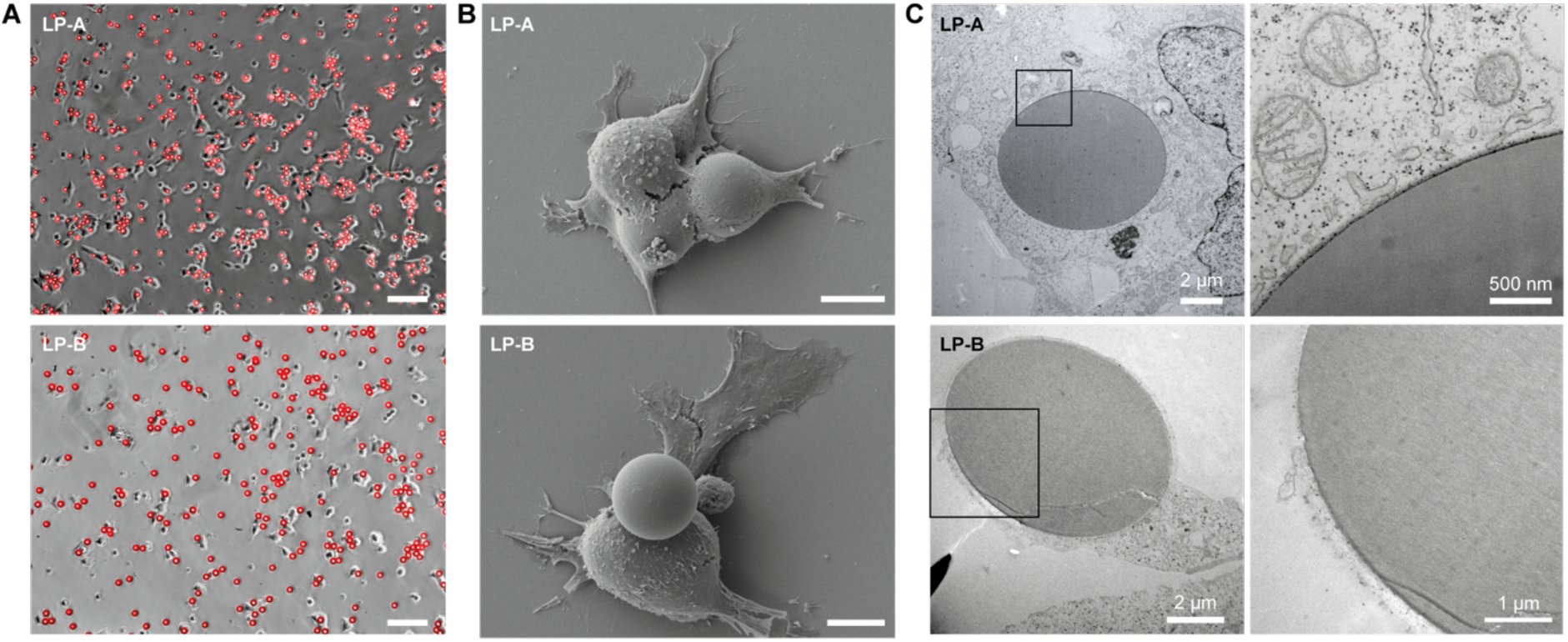
**A** Brightfield images of purified HEK293T cultures containing LP-A (top) and LP-B (bottom) at 24 h after cell sorting. Fluorescence of LPs is shown in red. Scale bar, 100 µm. **B** Representative SEM images of HEK293T containing LP-A (top) and LP-B (bottom) after cell sorting. Scale bar, 10 µm. **C** Representative TEM images of thin sections of HEK293T with LP-A (top) or LP-B (bottom). Squares indicate the zoomed-in regions shown on the right. The oval deformation of the LPs is an artefact of the sectioning process.

### Sorted populations of cells allow for control over LP-density in spheroids

To demonstrate that the flow cytometry based cell-sorting approach is suitable for high-throughput lasing experiments, cells with and without PEI-coated LPs were sorted into 96-well plates at defined ratios and cultured to form spheroids. In these optically dense spheroids, LPs were not visible in brightfield images (Fig. 4A) but could still be readily detected by their lasing signal using a hyperspectral confocal scanning microscope (Fig. 4B). The membrane-bound EGFP expression in HEK-EGFP enabled visualization of the spheroid by confocal fluorescence microscopy. Due to the precise z-plane control in confocal imaging, complete emission spectra of individual LPs at different positions within the approximately 300-µm-diameter spheroids could be recorded, allowing LP diameters to be calculated (e.g. 16.597 µm, 16.806 µm and 16.849 µm, Fig. 4C). Adjusting the relative populations of labeled and unlabeled cells yielded spheroids with defined numbers of LP-carrying cells. Round spheroids containing 10 or 20 LPs per 1000 cells formed within 36 h after seeding. Spheroids with higher numbers of LPs showed noticeable deformation, even after 36 h (SI Fig. 4). Lasing of LPs could be detected regardless of the number of LPs per spheroid and is displayed in a 3D hyperspectral reconstruction of the spheroid (Fig. 4D).

**Fig. 4.**
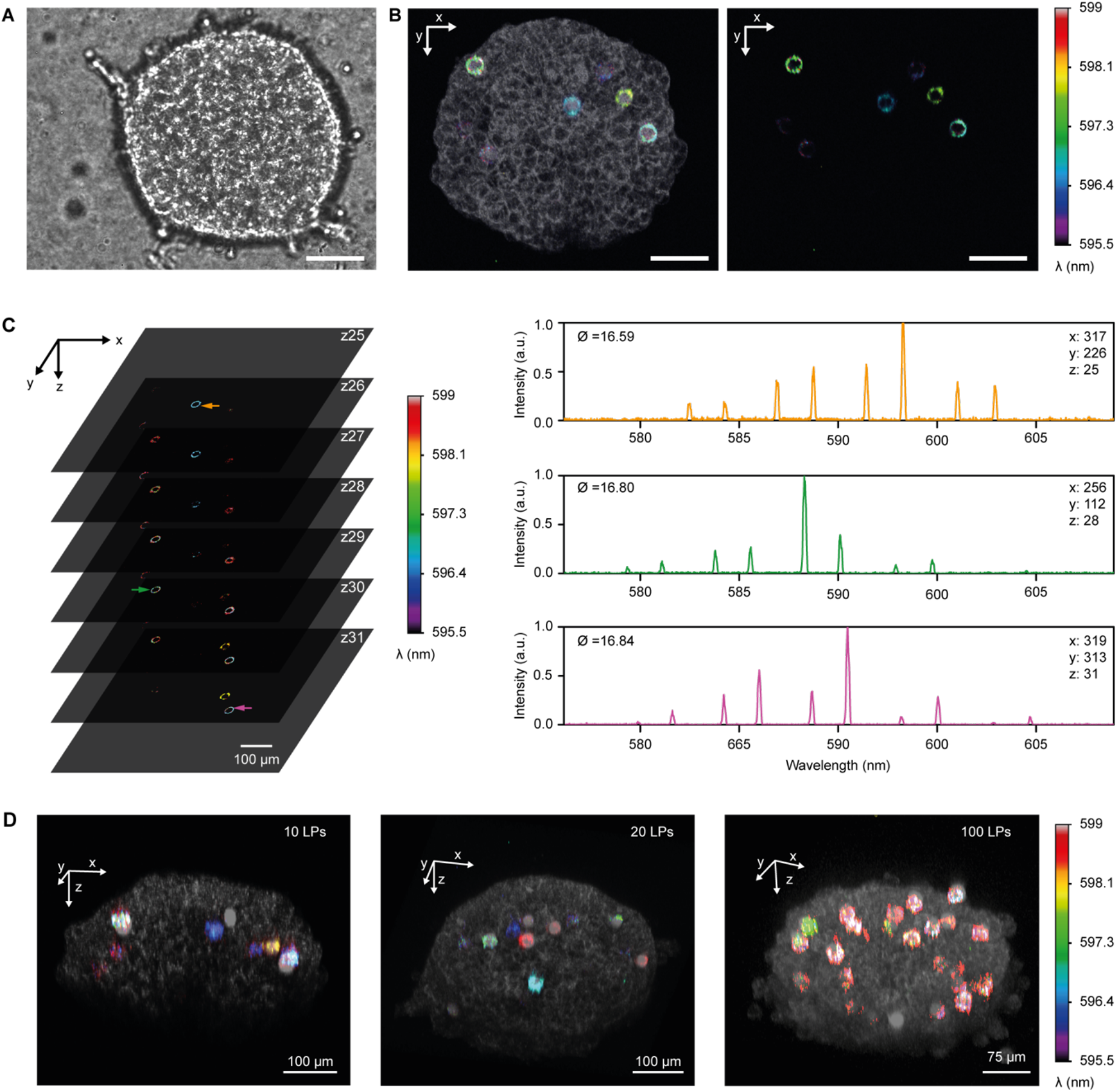
**A** Brightfield image of a HEK-EGFP spheroid containing LPs. Scale bar, 100 µm. **B** Maximum intensity projection of a hyperspectral confocal scan of a HEK-EGFP spheroid containing LP-B, with EGFP fluorescence shown in grayscale (left) and LP-B emission shown in a false-color scheme that encodes the wavelength of the most intense WGM lasing peak. Scale bar, 100 µm. **C** Confocal z-stack of the same spheroid as in B, with 5-µm spacing between each slice (left), and representative lasing spectra of the three LPs marked by arrows in the z-stack, taken from the hyperspectral confocal scan data. Note the high signal-to-noise ratio achieved despite measuring several hundred µm deep inside the tissue. **D** Three-dimensional representations of spheroids containing 10, 20 and 100 LPs at 24 h after FACS. EGFP fluorescence indicates cell membranes (grayscale). False-color scale indicates the wavelength of maximum lasing intensity.

## Discussion

In this study, we established and validated a workflow for sorting and analyzing cells labeled with large LPs using commercially available flow cytometry-based cell sorting and imaging flow cytometry platforms. Because the scattering properties of LPs and cells differ significantly, we are able to efficiently separate the different populations. The unique scattering properties of LPs in FACS are likely a result of the much higher refractive index of LPs compared to cells (*n* = 1.59 versus *n* = 1.4).

Beyond separating labelled from unlabeled cell populations, flow cytometry also proved to be a powerful tool for analyzing the size distribution, heterogeneity and fluorescence intensity of LPs. This revealed that lasing performance correlates strongly with the uniformity and fluorescence intensity of the LPs as measured by flow cytometry. LP-A exhibited a broader size distribution than LP-B and greater variability in lasing efficiency. The correlation between LP fluorescence intensity and SSC-A signal indicates that the greater fluorescence output mostly originates from the larger LP size, i.e., the larger volume LPs contains more dye molecules. Consequently, the broader size distribution of LP-A likely contributes to their lower absolute brightness.

The relatively large size and high refractive index of the LPs proved advantageous for separating LP containing from LP free cell populations based on forward- and side-scatter parameters in flow cytometry. However, differences in LP–cell interaction led to differences in sorting efficiency between the two LP types. The lower fraction of labelled cells in the LP-B sample compared with the LP-A sample may result from the partial attachment of LP-B to the cell surface, leading to a higher risk of detachment and subsequent loss during trypsinization and washing compared with the LP-A sample, where the LPs are mostly internalized. Control cell sorting experiments with LP-B alone revealed two subpopulations (SI Fig. 2C), suggesting potential aggregation which is not evident from microscopy. Furthermore, the variability observed in the LP^+^ fraction can be explained by multiple LPs attaching to individual cells, consistent with the heterogeneous attachment behavior seen in imaging experiments (SI Fig. 3C).

Besides the obvious difference in labelling efficiency between the two LP types detected by FACS, SEM and TEM, imaging also revealed that the two LP types interact very differently with cells. LP-A were fully internalized, with their surface in direct contact with the cytosol, whereas LP-B were at most partially engulfed, with the cell forming a membrane cup around them and their surface remaining closed off from the cytosol.

Earlier studies on polystyrene nanoparticles have shown that cellular uptake is most likely a two-stage process: initial electrostatic attachment to the cell surface, followed by an energy-driven endocytic mechanism^20^. During this initial electrostatic phase, the attachment strength of polystyrene microparticles has been found to correlate with their zeta potential, with more positively charged particles exhibiting stronger cell association, whereas neutrally charged particles show the weaker binding^19,21^. This is in line with our results, where the negatively charged bare LP-B did not attach to the cell surface. Coating the LPs with lipofectamine or PEI inverted the surface potential for both LPs to positive values but did not fully compensate for the differences in attachment or uptake efficiency. For instance, positively charged PEI–DNA polyplexes interact with negatively charged heparan sulfate proteoglycans on the cell membrane, illustrating how specific surface chemistries modulate cell-particle interaction ^22,23^. Surface groups of polystyrene LPs are determined by the fabrication processes involving manufacturer-specific initiators, surfactants, and catalysts, resulting in different surface chemistry and attachment behavior even in nominally identical LPs ^21^. Therefore, the use of polyvinyl alcohol in the synthesis of LP-B may contribute to reduced attachment to the cells ^24^.

While cells associated with either LP type were viable, the proliferation of cells with LP-B was reduced by approximately 20% after 5 days, indicating a potential inhibitory effect of these particles. Generally, exposure of cells to polystyrene nanoparticles is known to downregulate signaling cascades such as Akt, Erk, STAT5 and c-Src, leading to altered mitochondrial homeostasis and potentially impairing mitosis through ATP depletion^25^. Cells that have taken up polystyrene particles are also known to exhibit upregulated inflammatory genes, but these effects do not explain the difference in proliferation between samples with LP-A and LP-B^26^. However, PEI concentrations higher than 10 µg/ml are known to increasingly induce necrotic pathways and mitochondria-driven apoptosis^27^. The larger surface area of LP-B may lead to higher amounts of PEI bound to the surface, and therefore to higher PEI exposure to the cell.

Despite the somewhat negative effect of LP-B on cellular growth, their superior lasing performance will make them preferable for spheroid engineering in many cases. Overall, controlling the ratio of cells with and without LPs led to spheroids with the predicted number of LPs, even though some LPs detached during sorting or were damaged during centrifugation and therefore no longer lased when pumped. Lasing could be detected up to 300 µm deep inside the spheroids, consistent with previous studies^7^.

Commercial flow cytometry instruments have previously been used to distinguish cells with and without smaller associated polystyrene particles (4.5 to 6 µm in diameter) by analyzing particle fluorescence^28^. In immunology, cell sorting based on flow cytometry is also commonly applied to quantify particle uptake by phagocytic cells^29^. However, studies on the sorting of cells associated with particles large enough to exhibit lasing were still lacking in literature.

Lasing from 8 µm - 15 µm sized LP-A has previously been detected in a custom-built flow cytometry chip operated at a flow rate of 1.67 mm s^−^ (Ref. ^17^). In another study, small disk-shaped semiconductor LPs (2 µm in diameter) were used to label and sort cells at flow velocities of 500 mm s^−^ based on the lasing peaks of internalized LPs^16^. This approach has been extended to the simultaneous detection of 32 different antibodies, exceeding the capability of conventional flow cytometers. In addition, single-cell information was preserved through the identification of individual cells by the unique combination of LP lasing wavelengths^18^. Despite these advances, all reported approaches rely on custom-built microfluidic chips, complex optical setups, and advanced software for the detection and sorting of cells based on LP-associated lasing signals.

In contrast, our results demonstrate that commercially available flow cytometer and cell sorting systems can be readily adapted for the efficient purification and analysis of cell populations labeled with LPs using a purely scatter-based approach. This workflow provides an important foundation for advanced optical tagging strategies and the systematic investigation of cell–particle interactions.

In an age of increasing microplastic pollution, there is growing concern that LP-sized plastic microparticles present in the human environment can accumulate in the blood, potentially obstructing blood vessels in the brain, triggering inflammation, or exhibiting carcinogenic effects^10,30,31^. LPs may provide an attractive model to study these effects, given they can be readily tracked in complex environment and provide a read-out of changes in local density. Notably, LPs have recently been employed as an effective model for microplastic in marine organisms^12^. Our ability to label, sort, and monitor plastic microparticles provides a robust framework for systematically studying these interactions at the single-cell level and can thereby contribute to elucidating their broader physiological and ecological impact.

## Author Contributions

M.K. wrote the initial draft of the manuscript. A.P. and A.M.S. performed Zeta potential measurements. V.M.T. and T.T.P. performed hyperspectral imaging, T.T.P. calculated LP diameters and revised the manuscript. F.G. performed TEM imaging and guided the sample preparation for SEM. A.-L.S. and M.G. operated the cell sorter and imaging flow cytometer and revised the manuscript. M.S. and M.C.G. conceived and supervised the project and revised the manuscript.

## Acknowledgments

The authors thank Ruth Bruker for assisting with SEM. This work received financial support from the Humboldt Foundation (Alexander von Humboldt professorship) and the European Union’s Horizon 2020 Framework Programme (FP/2014-2020)/ERC grant agreement no. 640012 (ABLASE). This work received financial support from the European Research Council under the European Union’s Horizon Europe Framework Program/ERC Starting Grant agreement no. 101043047 (HYPERION, to M.S.). Instrument funding by the Deutsche Forschungsgemeinschaft in cooperation with the Ministerium für Kunst und Wissenschaft of North Rhine-Westphalia (INST 216/1120-1 FUGG, 469988234).

## Experimental Section

### Microscopy setup and lasing

The two LP types were obtained from different suppliers: LP-A from ThermoFisher (#36-4, Em_max_ = 612 nm) and LP-B from Microparticles GmbH (#PS-FluoRed-15.5, Em_max_ = 659 nm). LPs were characterized on a customized upright confocal Nikon Eclipse TE2000 microscope (Nikon Ltd., Japan) coupled to a SR500i spectrograph (Andor) according to established protocols^10^. For lasing comparisons of LPs from different suppliers, LPs were diluted in deionized water and left to dry on a coverslip. LPs were pumped at either 4 or 34 nJ with a pulsed laser (532 nm, 0.8 ns pulse width, 125 kHz repetition rate; Helios, Coherent, USA) through a Nikon Plan Apo 10x/0.45 objective (Nikon Ltd., Japan). For spheroid scanning, a water-dipping Nikon LWD 16x/0.8 was used.

### Surface modifications and measurement of surface potentials

LPs were resuspended in deionized water and subsequently washed with ethanol followed by three washes with deionized water. LPs were counted and their diameter measured on a Nexcelom Cellometer Auto T4 cell counter (Nexcelom, Bioscience Ltd., USA). Subsequently, the LPs were diluted to 1×10^5^ LPs/ml in deionized water, such that the total surface area across all samples was identical. 25 kDa PEI (#408727, Sigma Aldrich, USA) was dissolved to 1 mg/ml in deionized water, equilibrated to pH 7 with HCl, sterile filtered, and stored at -20°C. Either 10 µl of 1 mg/ml PEI or Lipofectamine 3000 (#L3000, Invitrogen, USA) were added per 100 µl LP solution. After 30 minutes incubation at room temperature, LPs were washed once in deionized water and either resuspended in DPBS (#14190250, Gibco, USA) for use in cells or in deionized water for surface potential measurements. Zeta potentials of all LPs were measured using a Malvern Zetasizer Nano ZS (Malvern Panalytical GmbH, Germany) to assess surface charge in aqueous suspension. Samples were dispersed in ultrapure water (resistivity 18.2 MΩ·cm at 25°C) and sonicated to ensure homogeneity. Measurements were performed at 25°C using plastic capillary cells (DTS1070). Electrophoretic mobility was determined via laser Doppler micro-electrophoresis and converted to zeta potential using the Smoluchowski approximation. Each outcome is an average of 5 scans, and results are reported as mean ± standard deviation.

### Uptake assay

HEK293T cells were cultivated at 37 °C and 5 % CO2, in DMEM (#21969035, Gibco, USA) with 10 % Fetal Bovine Serum (#A3160802, Gibco, USA), 1% Penicillin/Streptomycin (#15140-122, Gibco, USA) and 1 % GlutaMAX Supplement (#35050-038, Gibco, USA) and passaged at 80 % confluency. 24 h prior to the assay, cells were seeded at a density of 8×10^4^ cells per 35 mm gridded Ibidi dish (#81168, Ibidi, Germany). After 24 hrs media was changed to serum-free OptiMEM I (#31985062, Gibco, USA), LPs were added 30 minutes later. The next day, dishes were imaged and subsequently washed with DPBS and imaged again at the same position.

### Flow cytometry

Cells were seeded at a density of 1×10^6^ per well of a six-well plate. 24 h later the cell media was changed to OptiMEM I (#31985062, Gibco, USA) and LPs were added 30 minutes later. The next day, cells were dissociated for 60 s in 1 ml TrypLE Select (#12563029, Gibco, USA) and collected in FACS Buffer (DPBS, 2% FBS, 1% P/S). Cell sorting was performed using a BD FACSAria™ Fusion (with BD FACSDiva™ Software Version 8.0.1, BD Bioscience, San Jose, CA) software at RT using a 130 µm nozzle, 14 psi sheath pressure, and 0.9% NaCl as sheath fluid. Gating was based on FSC-A and SSC-A to identify three populations (bare LPs, HEK293T (LP^-^), HEK293T+LPs (LP^+^)). Cell aggregates and doublets were excluded using pulse-width parameters (FSC-W and SSC-W). Dead cells were excluded with DAPI (BV421-A, 405 nm excitation, 450/50 nm detection). Red photoluminescence from the LPs was detected in the PE-A channel. The Cytek Amnis ImageStream MkII Imaging Flow Cytometer was used for the validation of populations. Cells were sorted into Eppendorf tubes for further plating, cultivation, and imaging or into a 96 well plate for spheroid generation. Flow cytometry data was analyzed with FlowJo Software v10.10.0 (BD Life Sciences, Ashland, USA).

### Growth Assay

1.5×10^4^ HEK293T+LPs were seeded on a gridded imaging dish (#81168, Ibidi, Germany), and three fields of view (FOVs) were imaged after attachment. Confluency was automatically determined using Cellpose in Python^32^. The same FOVs were imaged every 2 h for 5 days. Results are reported as mean ± standard deviation.

### Electron Microscopy

For SEM, sorted HEK293T+LPs were plated on an indium tin oxide sputtered glass substrate inside a 3-well µ-dish insert (#80366, Ibidi) at a density of 1.5×10^4^/well, for TEM 1.5×10^4/^mm^2^ were seeded on Aclar and incubated overnight. For both, SEM and TEM, cells were fixed for 1 h in 2% glutaraldehyde (Sigma, #G5882-100ML) with 2.5 % sucrose (Roth, #4621.1) and 3 mM CaCl2 in 0.1M HEPES Buffer (Sigma-Aldrich, #C7902-500G) pH 7.4. Samples were washed three times with 0.1M HEPES Buffer and incubated with 1% osmium tetroxide (Science Services, #E19190) and 1% potassium hexacyanoferrate (Sigma-Aldrich, #P8131) for 1 h at 4°C. After 3x5min wash with 0.1M cacodylate Buffer (Applichem, #A2140,0100), samples were dehydrated at 4°C using ascending ethanol series (50%, 70%, 90%, 3x100%) for 7 min each. SEM samples were incubated with hexamethyldisilazane (HMDS) (Sigma-Aldrich, #440191-100ML) for 3 min. HMDS was removed and samples were dried under the hood. Samples were sputter coated with gold and imaged using a Zeiss Neon 40 Cross Beam operating at 5kV. TEM samples were infiltrated with a mixture of 50% Epon/ethanol for 1h, 70% Epon/ethanol for 2h and with pure Epon (Science Services, #E14120) overnight at 4°C. TAAB capsules (Agar Scientific, #G3744) filled with Epon were placed on top of the coverslip and cured for 48 h at 60°C. The coverslip was removed using liquid nitrogen. Ultrathin sections of 70 nm were cut using an ultramicrotome (Leica Microsystems, UC6) and a diamond knife (Diatome, Biel, Switzerland). Sections were stained with 1.5 % uranyl acetate (Agar Scientific, # R1260A) for 15 min at 37°C and with 3 % Reynolds lead citrate solution made from lead (II) nitrate (Roth, # HN32.1) and tri-sodium citrate dihydrate (Roth #4088.3) for 4 min. Images were acquired using a JEM-2100 Plus Transmission Electron Microscope (JEOL) operating at 80kV equipped with a OneView 4K camera (Gatan).

### Spheroid scanning

LP^+^ were sorted together with LP^-^ HEK293T cells in ratios of 1:10, 1:20 and 1:100 into wells of a ULA U-Bottom 96 well plate (#174952, ThermoFisher, USA) and incubated overnight. Spheroids were immobilized in 500 µl of 1.5 mg/ml Matrigel (#354230, Corning, USA) for 1 h at 37 °C. The Matrigel drop was fully covered with culture media. Spheroids were scanned on the hyperspectral confocal microscope setup, and data were analyzed using the previously published code^10^.

