## Supplemental Data for "Enriching microlaser-labelled cells by commercial flow cytometry-based cell sorting systems"

### Supplementary Information

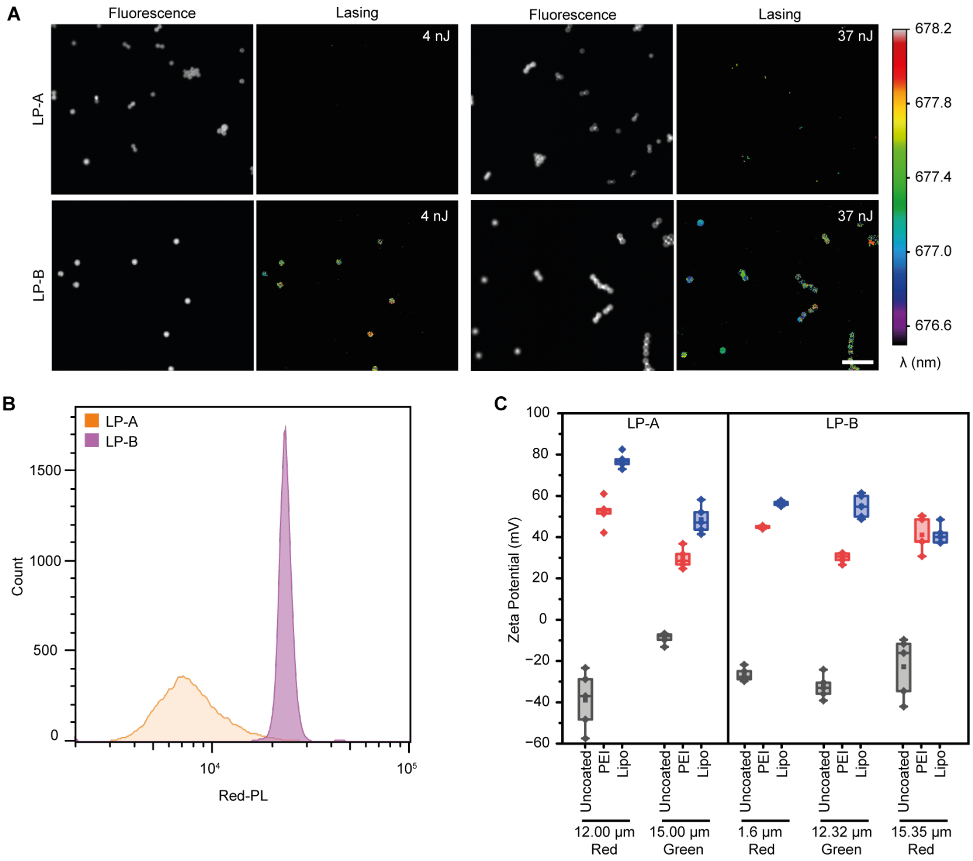


**SI Fig. 1**

**A** Fluorescence images (left) and corresponding lasing of LP-A and LP-B at two different pump energies, measured with a hyperspectral confocal microscope. Scale bar, 100 µm. At 4 nJ 100 % of LP-B and 7.22 ± 2.03 % of LP-A are lasing. At 34 nJ LP-A lasing increases to 40 ± 3.97 %.

**B** Fluorescence intensity (Red-PL) distribution in counts for LP-A and LP-B.

**C** Surface charge with and without coating for differently sized and dyed versions of the LP-A and LP-B particles. Line indicates median, square dot the mean, box shows 25-75% interval, whiskers depict 1.5 interquartile range, diamonds indicate outliers.

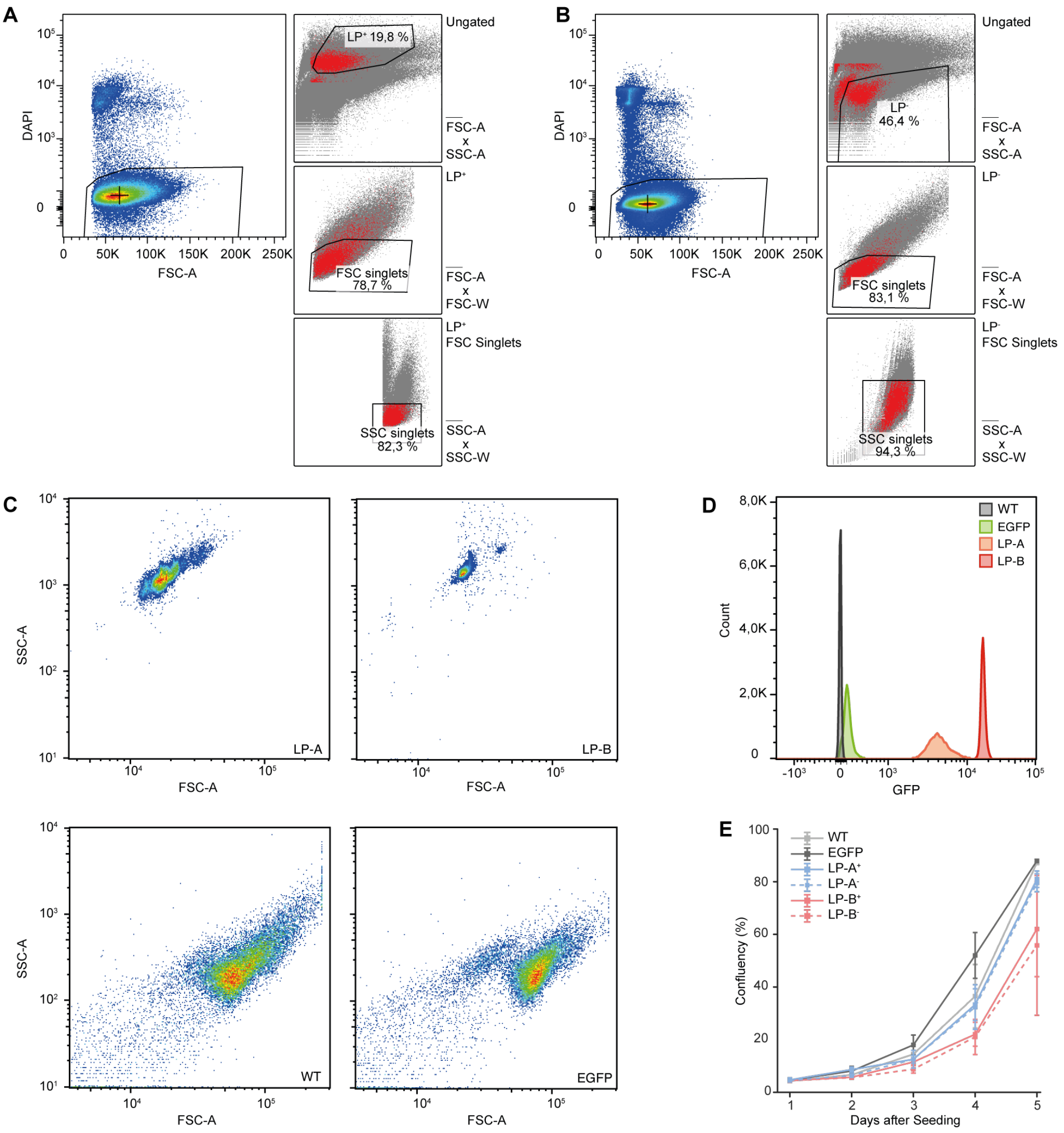


**SI Fig. 2**

**A** Backgating plots for the separation of viable LP⁺ from a HEK293T sample with LP-A (left). First LP⁺ were identified in FSC-A vs SSC-A (top right). The resulting population was subsequently gated for FSC singlets (center right) and SSC singlets (bottom right). LP⁺ singlets were finally sorted using DAPI intensity (left). FSC-A/W – Forward Scatter Area/Width, SSC-A/W – Sideward Scatter Area/Width.

**B** Same as A, but for the LP⁻ population of the same sample.

**C** FSC-A vs SSC-A properties of pure LP-A and LP-B as well as pure HEK-EGFP cells and their genetic background, wild-type HEK293T (WT).

**D** Fluorescence intensity comparison of WT, HEK-EGFP, and LPs in the FITC channel.

**E** Growth comparison of WT and HEK-EGFP, as well as HEK-EGFP exposd to LPs, after sorting, split into the groups LP⁺ and LP⁻.

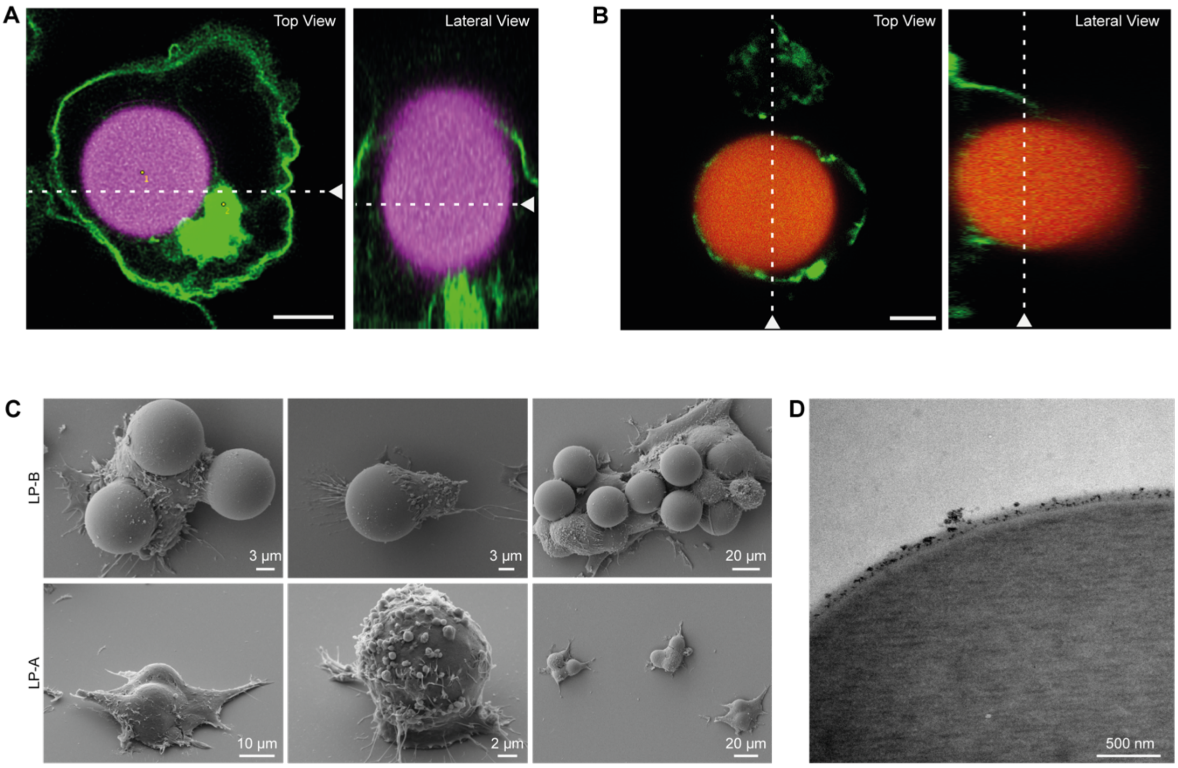


**SI Fig. 3**

**A** Confocal micrograph of a HEK293T cell containing an LP-A; horizontal (left) and vertical (right) planes are indicated with white arrows. Scale bar, 5 µm.

**B** Same as A for LP-B.

**C** SEM images of sorted HEK-EGFP cells associated with one or more PEI-coated LP-A or LP-B.

**D** LP-B surface in TEM with PEI aggregates on its surface.

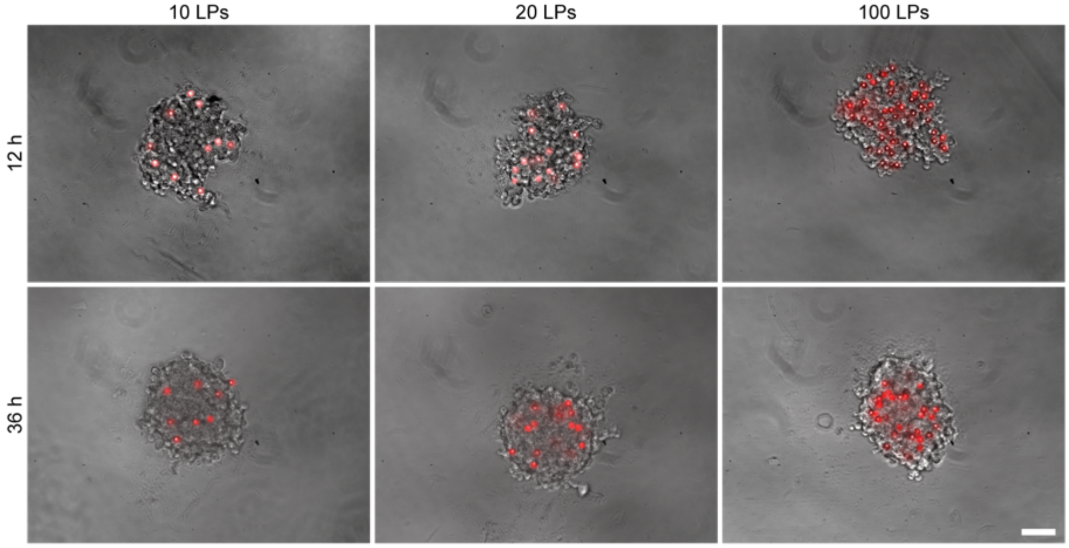


**SI Fig. 4**

Brightfield images of HEK293T spheroids with 10, 20 and 100 LPs per spheroid, 12 h (top) and 36 h (bottom) after sorting. LPs are indicated in red under fluorescence excitation.
